# Vitamin a deficiency causes apoptosis in the mammary gland of rats

**DOI:** 10.1101/2024.06.26.600922

**Authors:** M. Vasquez-Gomez, V. Filippa, M. Acosta, F. Mohamed, F. Campo Verde, C. Ferrari, G.A. Jahn, M.S. Giménez, D.C. Ramirez, S.E. Gomez Mejiba

**Affiliations:** Laboratory of Nutrition, Environment, and Metabolism. IMIBIO-SL. CONICET-San Luis, Universidad Nacional de San Luis. Av. Ejército de los Andes 950. San Luis, 5700 San Luis, Argentina; Laboratory of Histology. Universidad Nacional de San Luis. Av. Ejército de los Andes 950. San Luis, 5700 San Luis, Argentina; Laboratory of Reproduction and Breastfeeding. IMBECU. CONICET-Mendoza, Avenida Adrián Ruiz-Leal s/n, CC855, Mendoza, 5500 Mendoza, Argentina; Laboratory of Experimental and Translational Medicine. IMIBIO - SL. CONICET-San Luis, Universidad Nacional de San Luis. Av. Ejército de los Andes 950. San Luis, 5700 San Luis, Argentina; Laboratory of Nutrition and Experimental Therapeutics. IMIBIO-SL. CONICET-San Luis, Universidad Nacional de San Luis. Av. Ejercito de los Andes 950. San Luis, 5700 San Luis, Argentina

**Keywords:** Vitamin A deficiency, subchronic, mammary gland dysfunction, inflammation, apoptosis

## Abstract

Subchronic dietary vitamin A deficiency (VAD) causes severe abnormalities, including dysfunction of the epithelia of the mammary gland. However, the underlying mechanism of this process is partially known. Therefore, herein, we used a nulliparous-rat experimental model of dietary VAD for a total dieting regime of 3 and 6 months and intervened (refeed) with a vitamin A sufficient (VAS) diet for 0.5 or 1 month before the end of the diet regime and investigated the underlying molecular mechanism of mammary tissue dysfunction. Dietary vitamin A deficiency for 3 and 6 months caused increased inflammatory cell infiltration in the mammary gland parenchyma and glandular cells, with increased inflammation and apoptosis and reduced cell proliferation. These changes can be reversed with a VAS diet. Mammary gland dysfunction after a subchronic VAD is caused by an imbalance between NFκB and retinoic acid-triggered signaling. Inflammation, apoptosis, and impaired proliferation lead to dysfunction of the epithelia of the mammary gland of nulliparous rats fed a VAD diet.

## 1. INTRODUCTION

Worldwide, vitamin A deficiency (VAD) affects an estimated 190 million preschool-aged children and 19.1 million pregnant women^1^ ^2^, and it increases morbidity and mortality due to increased susceptibility to infection^3, 4^. Vitamin A and its derivatives (referred to as retinoids) are essential dietary compounds and are key regulators of cell differentiation, proliferation, inflammation, and death^1, 5–7^. Adult animals deprived of dietary vitamin A display severe abnormalities, including dysfunction of epithelia of the mammary gland, the mechanism of which is only partially known^1, 7^.

The mammary gland is a fascinating structural model for the study of apoptosis and its related signaling factors and underlying mechanisms ^5^. Its development is characterized by the stages of proliferation during puberty, pregnancy, and lactation, followed by an involution phase, when extensive tissue restructuring occurs, involving the removal of a proportion of secretory epithelial cells^7, 8^. In the context of the cell death program, the involution phase exhibits one of the most dramatic physiological responses, in which approximately 80% of mammary epithelial cells undergo programmed cell death^9^. One of the most important regulators of apoptosis via the mitochondrial pathway is the BCL2 protein family ^10^. The proapoptotic proteins BAX, BAK, and BAD, as well as the death suppressor proteins BCL-X, BCL2, and BCL-W, are synthesized in the mammary gland ^5^. Dynamic changes in the expression profiles of these proteins occur during development, suggesting that these changes may be involved in the regulation of postlactation apoptosis ^7, 11^. In vivo, cell survival programs can be determined by extracellular signals that directly affect the levels and function of members of the BCL2 family ^12^.

In mouse mammary gland tissue, BCL2 gene expression is undetectable during involution, whereas the BAX protein is detected in a large number of cells, and its expression increases during the first stage of involution^13–15^. Likewise, BAX mRNA levels increase with the onset of involution ^13^. On the other hand, in the mammary gland of BAX^-/-^ rats, involution is delayed at first but resembles the wild-type gland at 10 days after involution when remodeling is completed ^16, 17^. In a WAP-BAX transgenic mouse model of forced involution, it was demonstrated that the positive regulation of a single proapoptotic factor of the BCL2 protein family is sufficient to initiate the apoptosis of functionally differentiated mammary epithelial cells^18^. However, it is unknown whether VAD causes apoptosis in the mammary glands of virgin rats.

Retinoic acid (RA) interacts with nuclear receptors, such as the retinoic acid receptor (RAR) and retinoid receptor X (RXR), to regulate the transcription of several target genes by binding the retinoic acid-responsive elements (RAREs) in their regulatory region ^7, 19^. These receptors form heterodimers; RAR comprises three major isoforms (α, β, and γ) that interact with all forms of RA, whereas RXR, which also has the α, β, and γ isoforms, mainly interacts with *9-cis* RA. RA and its interaction with its cognate receptors RARα or RXR is critical for appropriate mammary gland differentiation and morphogenesis ^5, 19, 20^. RA- RARα/RXR interaction and binding to RARE has antioxidant and gene-transcriptional regulatory effects and plays a key role in cell differentiation, growth, proliferation, and immunity ^19, 21^.

The NFκB signaling pathway mediates important functions in cellular interactions, cell survival, and differentiation ^22^. These functions are mediated by the expression of several proinflammatory cytokines (e.g., TNFα and IL-6), enzymes (e.g., iNOS and COX-2), chemokines (e.g., CCL2 and CXCL10), adhesion molecules (e.g., ICAM-1 and VCAM-1) and proapoptotic genes, such as BAX ^23^. Vitamin A has an anti-inflammatory effect due to its binding to RARE, a consensus cis element found in several anti-inflammatory gene regulatory parts ^19^. Therefore, it is possible that under a VAD diet, the tight balance between NFκB and RA-activated RAR/RXR signaling is altered, in favor of NFκB signaling, in the mammary gland, causing tissue structural changes linked to apoptosis, proliferation, and cell differentiation.

Herein, we investigated whether VAD causes apoptosis in the mammary glands of nulliparous rats and investigated the underlying mechanisms by which this process occurs.

## 2. MATERIALS AND METHODS

### 2.1. Experimental model

Female Wistar rats bred in our animal facilities (IMIBIO-SL, National University of San Luis, Argentina) were used. Rats were weaned at 21 days old and immediately randomly assigned to 6 experimental groups. The experimental groups were as follows (see **Fig. 1** for a scheme of the experimental design): c3m, rats fed a VAS diet for 3 months; d3m, rats fed a VAD diet for 3 months; r3m, rats fed a VAD diet for 2.5 months followed by a VAS diet for

**Figure 1:**
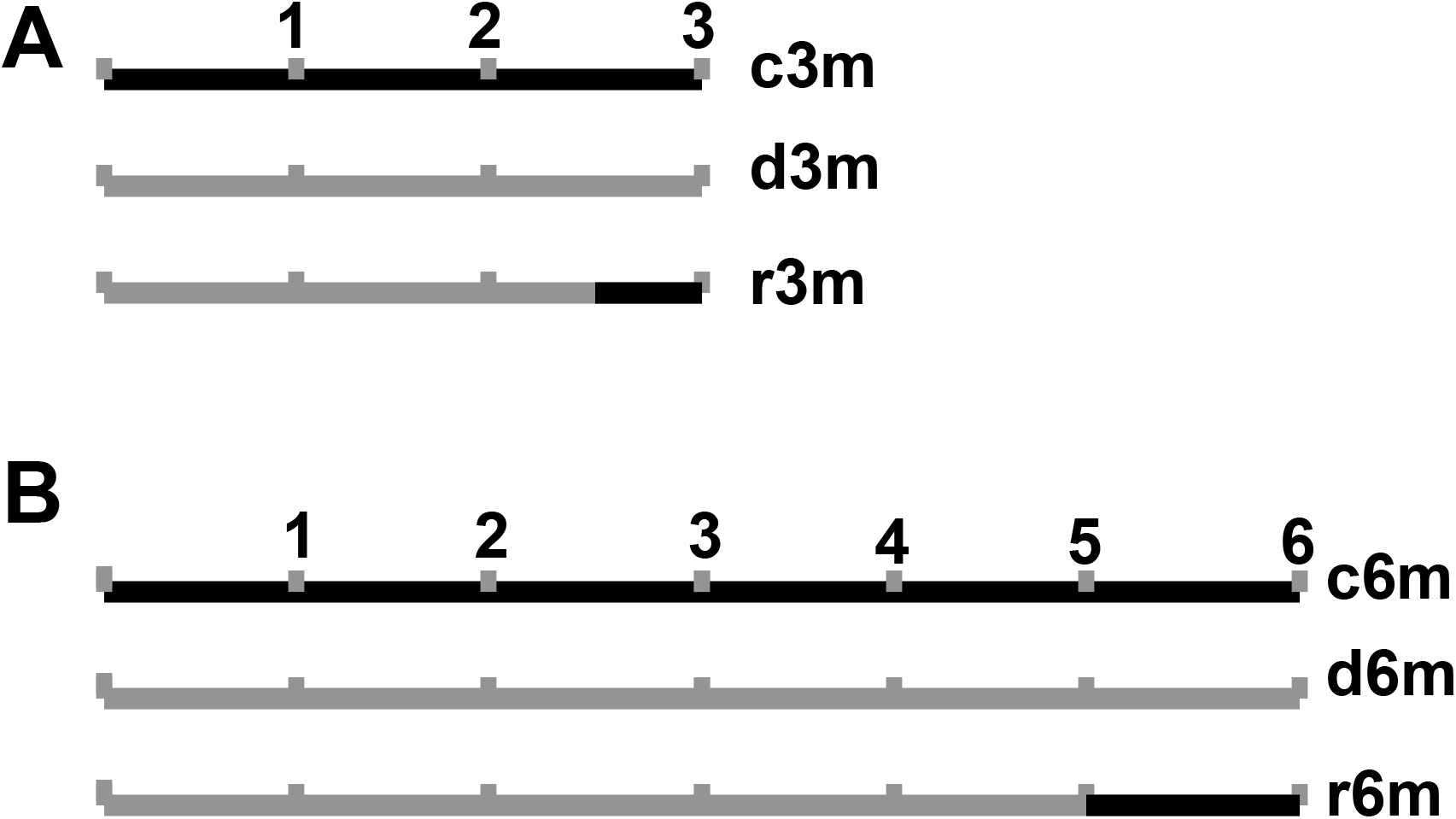
Experimental models: A) Three-month experimental models. B) Six-month experimental models. The black timeline indicates that rats were fed a VAS diet. The gray timeline indicates that rats were fed a VAD diet. Numbers in the timelines indicate months of diet. c3m, rats fed a VAS diet for 3 months; d3m, rats fed a VAD diet for 3 months; r3m, rats fed a VAD diet for 2.5 months, then fed a VAS diet for 0.5 months; c6m, rats fed a VAS diet for 6 months; d6m, rats were fed a VAD diet for 6 months; r6m, rats were fed a VAD diet for 5 months followed by a VAS diet for 1 month.

0.5 months; c6m, rats fed a VAS diet; d6m, rats fed a VAD diet for 6 months, r6m, rats fed a VAD diet for 5 months followed by a VAS diet for 1 month. Diets were prepared according to the AIN-93G for laboratory rodents ^24^. The composition of the VAD and VAS diets is shown in **Table S1**, and the composition of the mineral and vitamin mixtures is shown in **Table S2** and **Table S3**, respectively. The only difference between the VAD and VAS diets is that the vitamin mixture added to the VAD does not contain trans-retinyl palmitate. The rats were housed in individual cages and kept in a 21–23 °C controlled environment with a 12-hour light:dark cycle. They were given free access to food and water throughout the entire 3 and 6 months of dieting. After the entire treatment period, the rats were euthanized by CO_2_ inhalation. The whole experiment with animals was performed according to the National Institutes of Health Guide for the Care and Use of Laboratory Animals^25^ and the National University of San Luis Committee’s Guidelines for the Care and Use of Experimental Animals (Approved UNSL-CICUAL-Protocol# B-202-3/15-FQByF-RD-324- 16).

### 2.2. Measurement of retinol concentrations in the plasma and mammary gland

The blood samples were collected in EDTA-treated tubes. To minimize photoisomerization of vitamin A, the plasma was taken under reduced yellow light and frozen in the dark at –70°C until the determination of retinol concentrations. Analyses were carried out within 1–3 weeks of obtaining the samples. The retinol concentration was determined by the modified technique of Neeld and Pearson^26^. Briefly, the plasma was treated with 1 ml of 95% ethanol to precipitate proteins. The supernatant was then treated with 1.5 ml of petroleum ether to extract vitamin A and carotenoids. After centrifugation at 3000 rpm for 10 minutes, the supernatant was read at 450 nm, corresponding to the β-carotene absorbance, and then dried in an oven at 37 °C. The residue was taken up in 50 µl of chloroform, 50 µl of acetic anhydride, and 500 µl of trifluoroacetic acid with vigorous stirring. The samples were read within 30 seconds at 620 nm. A standard curve of vitamin A and β-carotenes was processed. Because β-carotenes react with trifluoroacetic acid, the results were corrected after reading the absorbance at 450 nm and calculating the corresponding correction factor.

### 2.3. Histology of the mammary gland

For the light microscopy studies, four mammary glands from each group were excised and fixed in Bouin’s fluid. The samples were then dehydrated in an increasing ethanol series, cleared in xylene, and embedded in paraffin. Sections of 5 μm thickness were obtained using a Reichert-Jung Hn 40 sliding microtome. In these sections, hematoxylin & eosin (H&E) staining, immunohistochemistry, and TUNEL assays were performed.

### 2.4. Immunohistochemistry

BCL2- and BAX-positive cells were detected by immunohistochemistry in paraffin slides of the mammary gland. Briefly, the sections were first deparaffinized with xylene, hydrated through decreasing concentrations of ethanol, and rinsed with distilled water and phosphate-buffered saline (PBS, 0.01 M, pH 7.4). Antigen retrieval was performed by microwaving the sections for 6 min (2 x 3 min) at full power in sodium citrate buffer (0.01 M, pH 6.0). Endogenous peroxidase activity was inhibited with 3% H_2_O_2_ in water for 20 min. Nonspecific binding sites for immunoglobulins were blocked by incubation for 20 min with normal mouse serum diluted in PBS containing 1% bovine serum albumin, 0.09% sodium azide, and 0.1% Tween-20. Sections were incubated with the following primary antibodies: 8 hours in a humidified chamber at 4 °C with mouse monoclonal anti-human BCL-2 (clon BCL-2/100; Catalog. No. AM287-5 M, BioGenex, San Ramón, CA, USA), 30 min in a humidified chamber at 20 °C with rabbit polyclonal anti-human BAX protein (Catalog. No. AR347-5R, BioGenex, San Ramón, CA, USA), and 4 hours in a humidified chamber at 20 °C with mouse monoclonal anti-rat proliferating cell nuclear antigen (PCNA) (clon PC10; Catalog. No. AM252-5 M, BioGenex, San Ramón, CA, USA). After rinsing with PBS for 10 min, immunohistochemical visualization was carried out using the Super Sensitive Ready-to-Use Immunostaining Kit (BioGenex, San Ramon, CA, USA), which was used as follows: sections were incubated for 30 min with diluted biotinylated anti-IgG and, after being washed in PBS, were incubated for 30 min with horseradish peroxidase-conjugated streptavidin and finally washed in PBS. The reaction sites were visualized using a freshly prepared solution containing 100 µL of 3,3’-diaminobenzidine tetrahydrochloride chromogen in 2.5 mL of PBS and 50 µL of H_2_O_2_ substrate solution. The sections were counterstained with Harris’ hematoxylin for 10 s, dehydrated, and mounted. For the negative control of immunohistochemistry, the following procedure was performed: 10% normal serum and PBS replaced the primary antibodies. No positive structures or cells were found in these sections. The large and small intestines of rats were used as positive controls.

### 2.5. TUNEL assay

The Terminal deoxynucleotidyl transferase (TdT)-mediated biotinylated dUTP Nick-End Labeling (TUNEL) assay [Dead-End Colorimetric TUNEL System, Promega, Madison, WI] was used according to the manufacturer’s instructions. A positive control was run after pretreatment with DNAse I. A negative control was produced by using the equilibrium buffer mixture with biotinylated nucleotides but without the TdT enzyme. Slides were counterstained with hematoxylin and mounted without contrast agents.

### 2.6. Morphometric analysis of immunohistochemistry images

The mammary gland slides used for assessing BCL2-, BAX-, TUNEL-, and PCNA-positive cells were examined using an Olympus BX-40 light microscope. Immunohistochemical images were captured by a Sony SSC-DC5OA camera and processed with Image-Pro Plus 5.0 software. Briefly, the image was displayed on a color monitor; a standard area of 18141.82 µm^2^ (reference area) was defined on the monitor, and distance calibration was performed using a slide with a micrometric scale for microscopy (Reichert, Austria). The morphometric study was performed as follows: three regularly spaced serial tissue sections (100 µm each) from mammary glands were used, and microscopic fields were examined under 400× magnification. In each section, 10 microscopic fields were randomly selected throughout the mammary gland. In each image, the percentages of TUNEL- and PCNA- positive cells were determined using the following formula: Percent of positive cells = positive cells/Total cell count×100.

BAX^+^ and BCL2^+^ cells were measured using an Image Pro-Plus 3.0.1 system (Media Cybernetics, Silver Spring, MA, USA). For the immunohistochemistry technique, images were digitized by a color charge-coupled device (CCD) video camera (Sony, Montvale, NJ, USA) mounted on a conventional light microscope (Olympus BH-2; Olympus Company, Tokyo, Japan) using a magnification of 400x. The microscope was prepared for Koehler illumination. This was achieved by recording a reference image of an empty field for the correction of unequal illumination (shading correction) and by calibrating the measurement system with a reference slide to determine background threshold values. The reference slides contained a series of tissue sections stained in the absence of a primary antibody. The positive controls were used as interassay controls to maximize the levels of accuracy and robustness of the method. The methodological details of image analysis have been described earlier^27^. Using a color segmentation analysis tool, the total intensity of the positively stained cytoplasmic area (brown reaction product) was measured and is shown as a ratio (%) of the total cytoplasmic area (brown reaction product blue hematoxylin). The image analysis score was calculated separately by using AutoPro macro language, an automated sequence operation created to measure the immunohistochemical-stained area (IHCSA). The IHCSA was calculated as a percentage of the total area evaluated through color segmentation analysis, which extracts objects by locating all objects of a specific color (brown stain). The brown stain was selected with a sensibility of 4 (maximum 5), and a mask was then applied to separate the colors permanently. The images were then transformed to a bilevel scale Tagged Image File Format (TIFF). The IHCSA (black area) was calculated from at least fifty images of each area of the mammary gland in each slide being studied.

### 2.7. RNA isolation and RT**L_**qPCR analysis

The mammary tissues were homogenized with the Ultra-Turrax T-25 digital homogenizer. Total RNA was isolated from 150-200 mg of mammary tissue using the guanidinium isothiocyanate-acid phenol method as modified by Puissant and Houdebine^28^. Ten micrograms of total RNA was reverse transcribed (RT) at 37 °C using random hexamer primers and Moloney murine leukemia virus retrotranscriptase (Invitrogen-Life Technologies, Buenos Aires, Argentina) in a 20 µL reaction mixture. The RNA was first denatured at 70 °C for 5 min in the presence of 2.5 µg of random hexamer primers (Invitrogen). For the subsequent RT reaction, the following mixture was added: RT buffer [50 mM Tris-HCl (pH 8.4), 75 mM KCl, 3 mM MgCl_2_], 0.5 mM dNTPs, 5 mM DTT, 200 units M-MLV Reverse Transcriptase. The reaction was incubated at 37 °C for 50 min. Next, the reaction was inactivated by heating at 70 °C for 15 min. The cDNA was stored at -20 °C. The mRNA levels of BAX, BCL-2, and S16 were estimated by real-time RT[PCR using the rat-specific primers and reaction conditions described in supplementary **Table S4**. The PCRs were performed using a Corbett Rotor-Gene 6000 Real-Time Thermocycler (Corbett Research Pty Ltd. Sydney, Australia) and Eva-Green™ (Biotium Hayward, CA) in a final volume of 20 µL. The reaction mixture consisted of 2 µL of 10X (PCR Buffer, 1 µL of 50 mM MgCl_2_, 0.4 µL of 10 mM dNTP Mix (Invitrogen), 1 µL of Eva Green, 0.25 µL of 5 U/mL Taq DNA Polymerase (Invitrogen), 0.1 µL of each 2.5 mM primer (forward and reverse primers) and 10 µL of diluted cDNA. The PCRs were initiated with incubation for 5 min at 95 °C, followed by 40 cycles. Melting curve analysis was used to check that a single specific amplified product was generated. Real-time quantification was monitored by measuring the increase in fluorescence caused by the binding of EvaGreen™ dye to double-strand DNA at the end of each amplification cycle. Relative expression was determined using the comparative quantitation method of normalized samples with the expression of a calibrator sample, according to the manufacturer’s protocol ^29^. Each PCR run included a no-template control and a sample without RT. All measurements were performed in duplicate. The reaction conditions and quantities of cDNA added were calibrated so that the assay response was linear for the amount of input cDNA for each pair of primers. RNA samples were assessed for DNA contamination by performing different PCRs without prior RT. Relative levels of mRNA were normalized to the S16 or S28 housekeeping gene expression. The real-time PCR products were analyzed on 2% agarose gels containing 0.5 mg/mL ethidium bromide, and a unique band of approximately correct molecular weight corresponded with a unique peak in the melt curve analysis.

### 2.8. Statistics

The normality of the data was determined with the Shapiro-Wilk test. The data are shown as a representative image or mean values ± standard deviations (SD) and were analyzed by two-way ANOVA followed by Tukey’s *post hoc* test for specific comparisons. p<0.05 was considered to be significant. GraphPad Prism (v. 3.02) software was used for statistical analysis.

## 3. RESULTS

### 3.1. Dietary VAD affects body weight gain in virgin rats

Body weight gains after 3 or 6 months of diet were determined in each experimental group (**Table 1**). After 3 or 6 months of diet, animals fed a VAD diet gained less weight than those fed a VAS diet (c3m vs d3m and c6m vs d6m), with a statistically significant difference (p<0.05). These changes were reversed in the r3m and r6m groups, with a statistically significant difference (p<0.05) between the r3m vs d3m and r6m vs d6m groups.

**TABLE 1:**
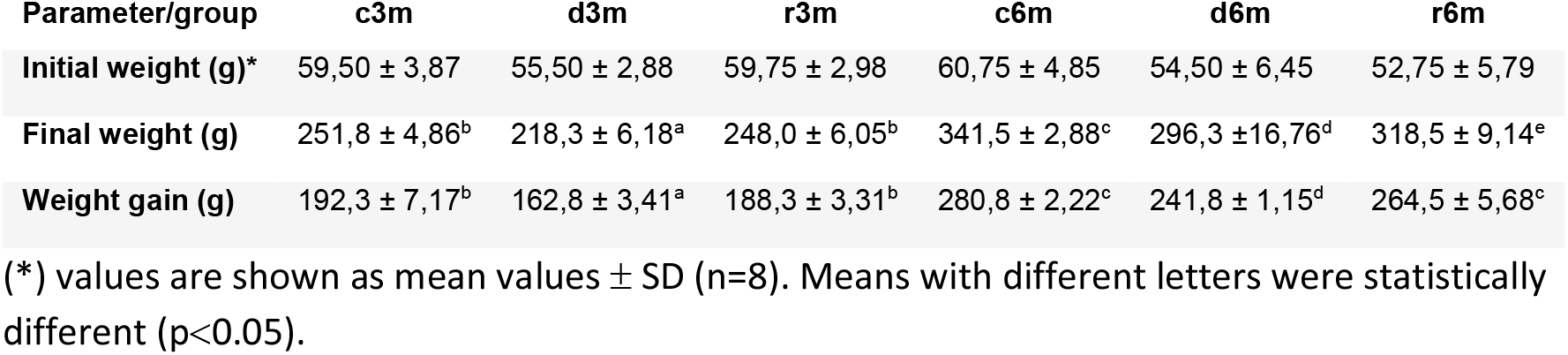
Body weight gain in the different experimental groups.

### 3.2. Dietary VAD affects the histological architecture of the mammary gland

To determine the effect of VAD on the tissue architecture of the mammary gland, we performed a morphological study in H&E-stained slides (**Fig. 2)**. In the mammary glands of the c3m rats, ductal development was observed; however, it was not observed in the d3m rats. In the mammary gland of the c3m group, there were more ducts than in the deficient group, and there was a beginning of incipient alveolar lobe arborization. In the r3m rats, modest alveolar lobe development is observed, greater than in d3m rats. In the d6m group, lower parenchymal development (i.e., fewer acini or alveoli) was observed, highlighting the ductal predominance compared to the c6m group. In the animals from the r6m group, the recovery of the size and number of mammary alveoli stands out, similar to that observed in the c6m group. A subchronic deficiency of dietary vitamin A causes mammary gland dysfunction in the mammary gland of nulliparous rats. These findings are consistent with an abnormal development of the mammary gland parenchyma. Dietary vitamin A supplementation prevents mammary gland dysfunction in nulliparous rats.

**Figure 2:**
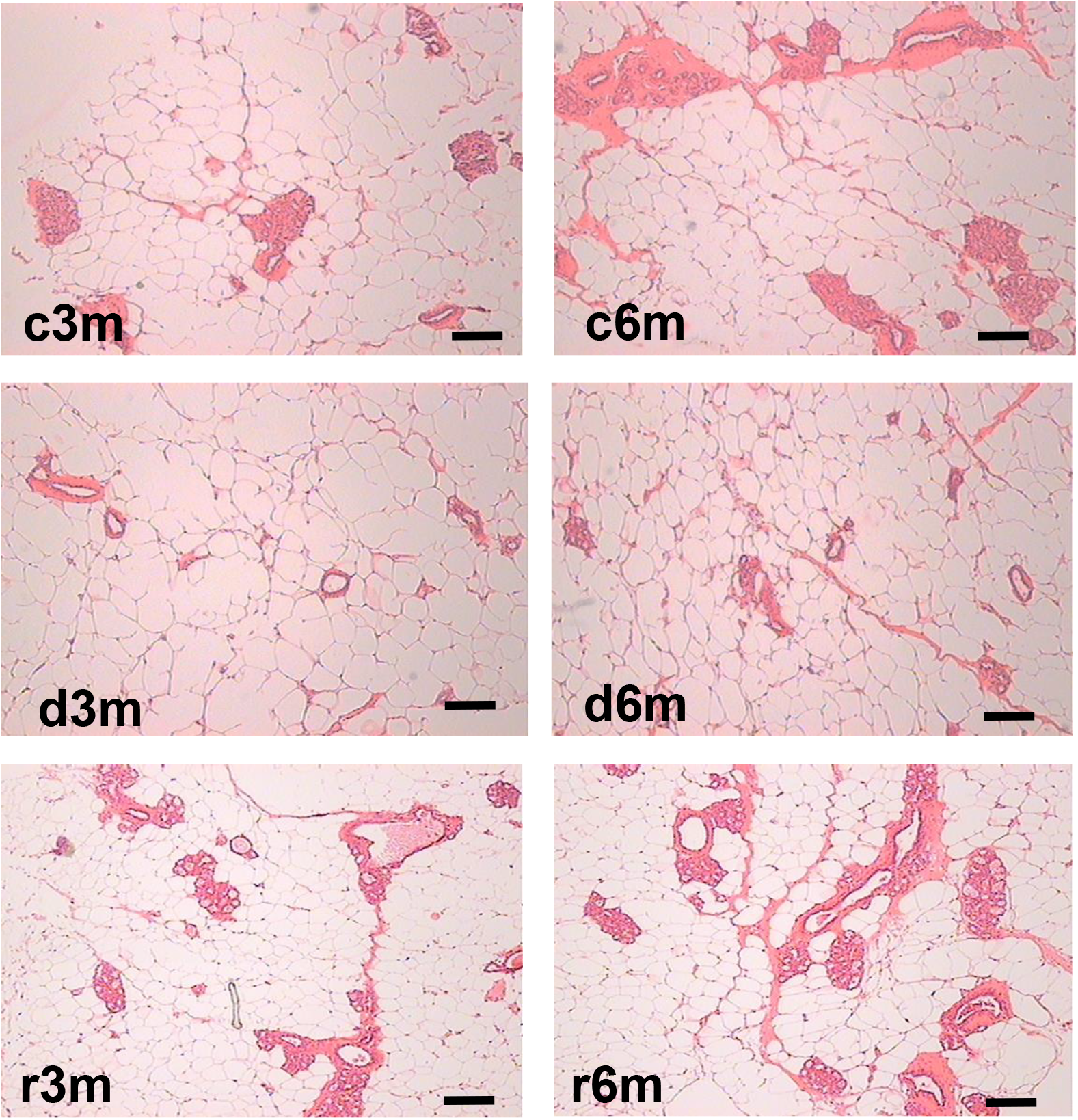
Mammary gland structural changes caused by VAD. H&E staining of mammary glands isolated from the experimental groups c3m, d3m, r3m, c6m, d6m. Magnification 40X. The scale bar is 100 µm.

### 3.3. Dietary VAD alters RA concentration in serum, liver, and mammary gland

The effect of dieting on the concentration of RA in the serum, liver, and mammary gland in animals from each experimental group (see Fig. 1) is shown in **Table 2**. A VAD diet for 3 months (d3m) caused a reduction in the RA concentration in serum, liver, and mammary glands. Although refeeding animals for 15 days (r3m) only partially reestablished (one-third) the concentration of RA found in those tissues from animals fed a VAS diet for 3 months (c3m). The concentrations of RA in the serum, liver, and mammary gland of animals fed a VAS diet for 3 or 6 months were similar (c3m vs c6m). However, RA concentrations in the serum of animals fed a VAD diet for 6 months were approximately 30 times lower than those fed the same diet for 3 months (d6m vs d3m). The effect of refeeding a VAS diet for 1 month increased RA serum and liver concentrations similar to those of feeding a VAS diet for 6 months (r6m vs c6m). However, the concentration of RA in the mammary gland increased, but only approximately one-third of those found in the mammary gland of rats fed a VAS for 6 months (r6m vs c6m).

**TABLE 2:**
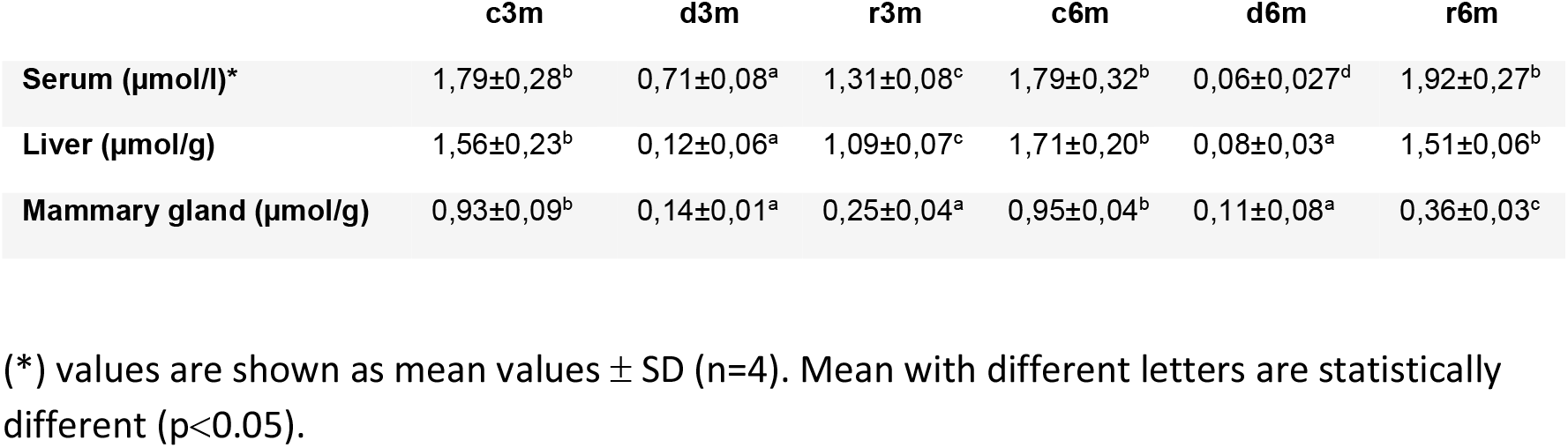
Retinoic acid concentrations in serum, liver, and mammary gland.

|  | c3m | d3m | r3m | c6m | d6m | r6m |
| --- | --- | --- | --- | --- | --- | --- |
| **Serum (µmol/l)\*** | 1,79±0,28<sup>b</sup> | 0,71±0,08<sup>a</sup> | 1,31±0,08<sup>c</sup> | 1,79±0,32<sup>b</sup> | 0,06±0,027<sup>d</sup> | 1,92±0,27<sup>b</sup> |
| **Liver (µmol/g)** | 1,56±0,23<sup>b</sup> | 0,12±0,06<sup>a</sup> | 1,09±0,07<sup>c</sup> | 1,71±0,20<sup>b</sup> | 0,08±0,03<sup>a</sup> | 1,51±0,06<sup>b</sup> |
| **Mammary gland (µmol/g)** | 0,93±0,09<sup>b</sup> | 0,14±0,01<sup>a</sup> | 0,25±0,04<sup>a</sup> | 0,95±0,04<sup>b</sup> | 0,11±0,08<sup>a</sup> | 0,36±0,03<sup>c</sup> |

(\*) values are shown as mean values ± SD (n=4). Mean with different letters are statistically different (p<0.05).

### 3.4. Dietary VAD causes apoptosis in the mammary gland of virgin rats

Figure 3A shows the results of BAX immunohistochemistry in the mammary glands of the different experimental groups. An increase in BAX immunostaining was observed in the deficient groups of 3 and 6 months for the control and refed groups, respectively. This is corroborated by cellular quantification (Fig. 3B). The percentage of BAX+ cells is increased in the vitamin A-deficient groups relative to the respective controls and refed. The observed increments were only reversed after vitamin A refed for 1 month in the deprived groups for 5 months. We then corroborated the immunohistochemistry data by measuring BAX gene expression relative to ribosomal S16 gene expression. As observed in **Fig 3C**, BAX gene expression follows a similar pattern as observed in BAX+ cell counts shown in 3A.

**Figure 3.**
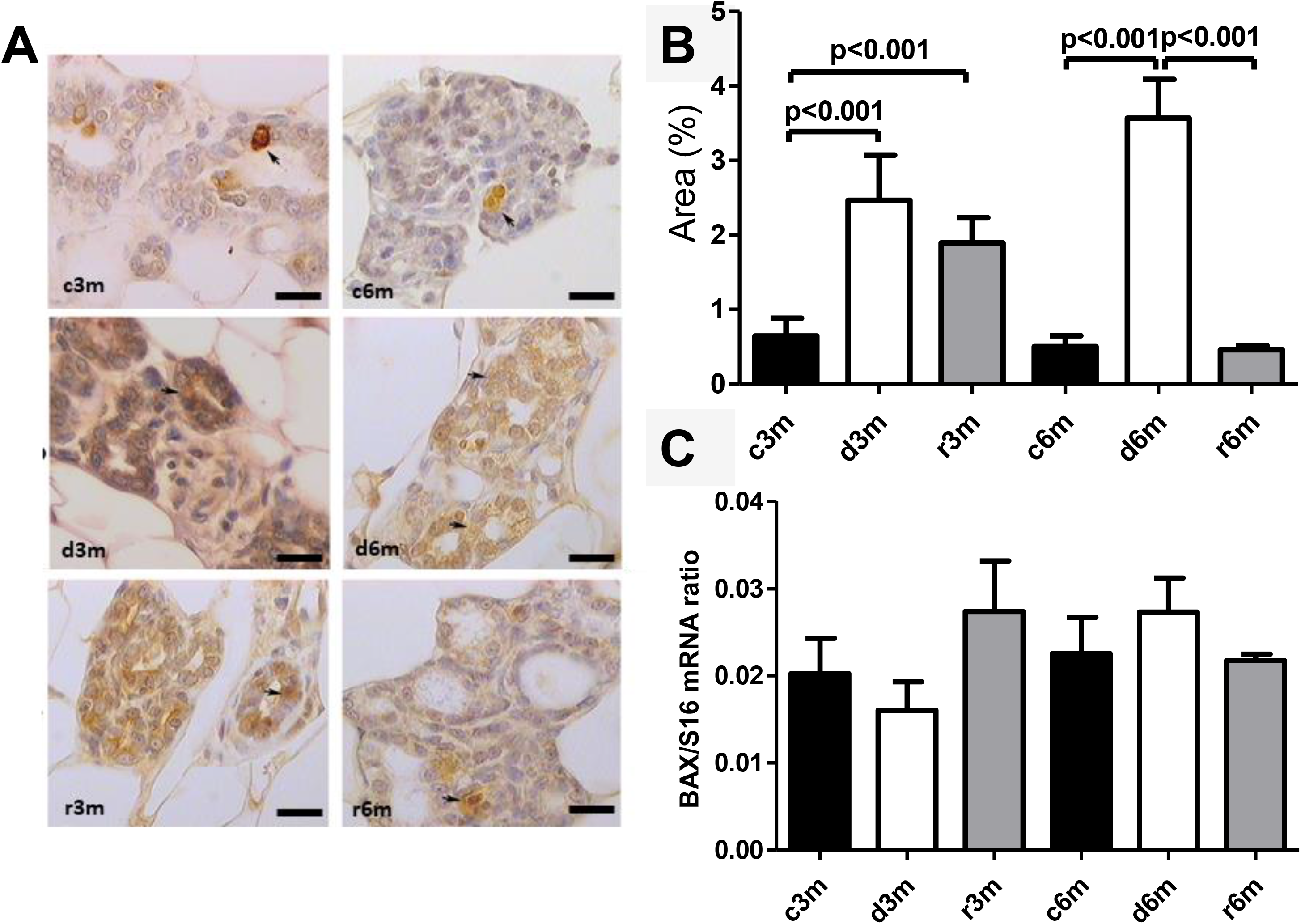
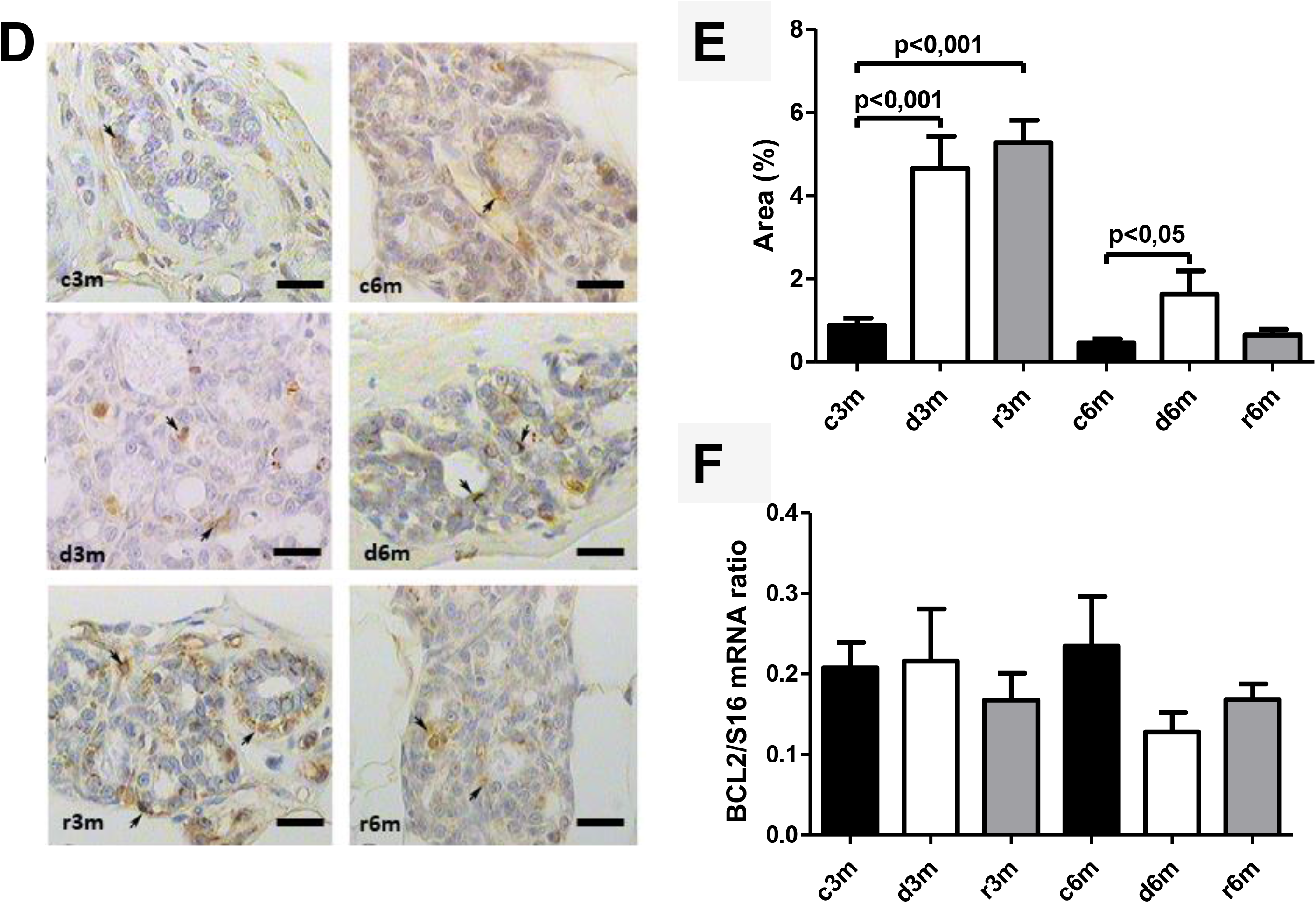
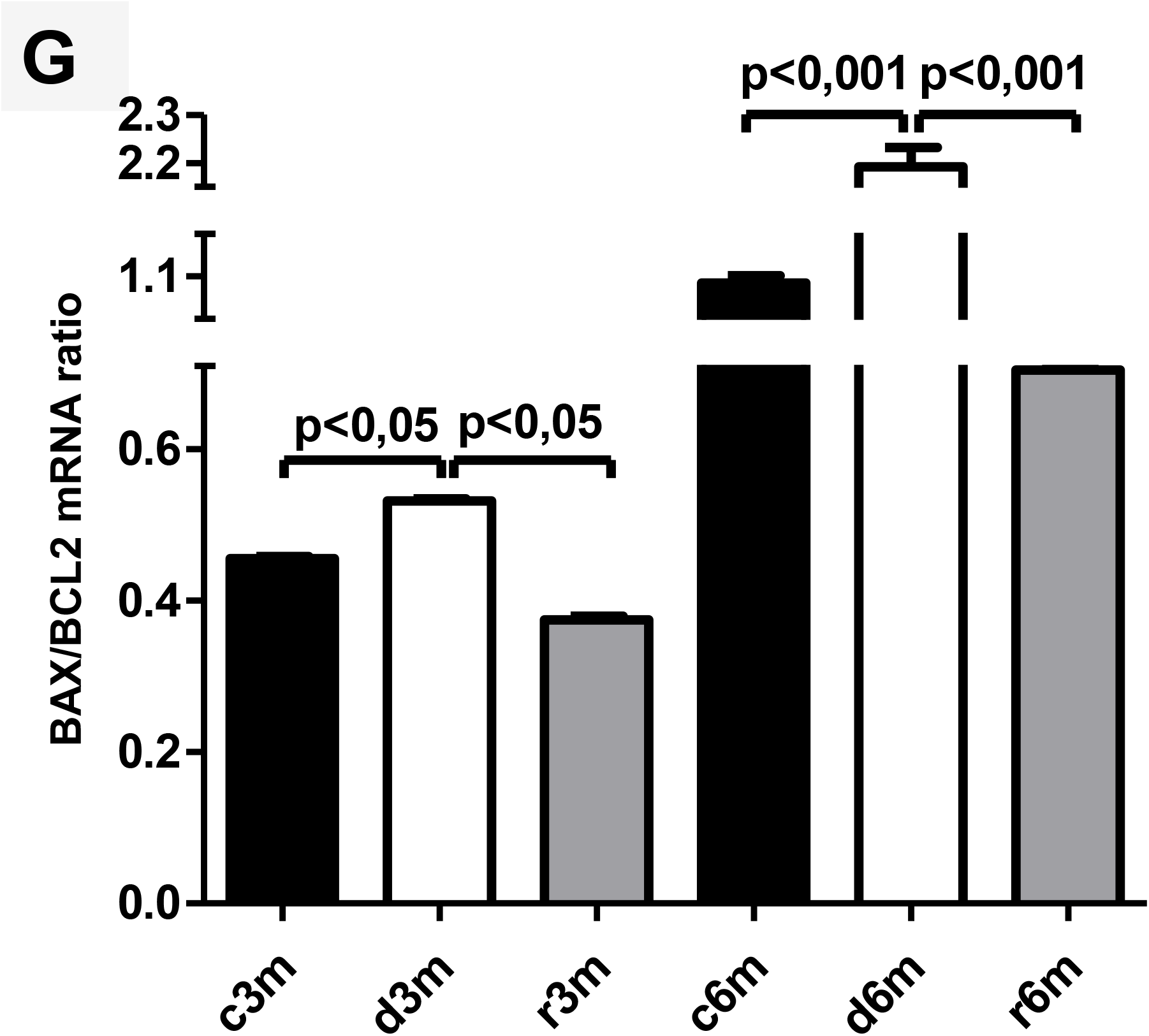
VAD and further refeeding with a VAS diet cause changes in BAX and BCL2 gene expression in the mammary gland. A) Anti-BAX immunohistochemistry images in the mammary gland. Black arrows indicate positive immunostaining. Magnification 100×. Scale bar: 25 µm. B) Percentage of positive area for anti-BAX immunohistochemistry images shown in A. C) Measurement of bax gene expression as assessed by RT[qPCR in the mammary gland. D) Anti-BCL2 immunohistochemistry images in the mammary gland. Magnification 100×. Scale bar: 25 µm. E) Percentage of positive area for anti-BCL2 immunohistochemistry images shown in A. F) Measurement of BCL2 gene expression as assessed by RT[qPCR. G) BAX/BCL2 gene expression ratio from immunohistochemical images shown in A and D. In A and D, the scale bar is 25 µm, and black arrows indicate positive cells. BAX and BCL2 gene expression levels are shown relative to the expression of the constitutive ribosomal S16 gene. The data shown are representative images or mean values ± SD (n = 4).

Figure 3D shows the results of BCL2 immunohistochemistry in the mammary gland of the different experimental lots, where an increase in BCL2 immunolabeling was observed in the deficient group at 3 months (d3m) compared to the control group (c3m). Refed with vitamin A for 3 months does not revert to the values of the controls. In 6-month control rats (c6m), little cytoplasmic immunostaining is observed. Feeding rats for 6 months a VAD diet caused increased BCL2 immunostaining compared to those rats fed a VAS diet for 6 months (d6m vs c6m). These observations were corroborated by quantification of BCL2+ (Fig. 3E). BCL2+ cells in the mammary gland increased in the deficient lots at 3 and 6 months relative to the respective controls. **Fig 3F** shows BCL2 gene expression in the mammary glands of the different experimental groups as assessed by RT[qPCR. Figure 3G shows the BAX/BCL2 gene expression ratios as assessed by measuring gene expression by RT[qPCR. In the 3- and 6-month dietary regimes, dietary vitamin A deficiency increased the BAX/BCL2 gene expression ratio in the mammary glands. Refeeding with a VAS diet for 0.5 or 1 month restored the BAX/BCL2 gene expression ratio in the mammary gland. Dietary vitamin A prevents apoptosis in the mammary gland of nulliparous rats.

### 3.5. Dietary deficiency of vitamin A increases apoptosis in the mammary gland parenchyma

To corroborate the immunohistochemistry and gene expression analysis of molecular markers of apoptosis (see Fig. 3), we performed a TUNEL assay (Fig. 4A). In the mammary gland of the 3-month control group, there were few parenchymal cells with apoptotic nuclei. In the d3m group, numerous glandular cells with apoptotic nuclei were observed, evidencing a remarkable increase with respect to the control group (d3m vs c3m). These data are consistent with the diminution of glandular parenchyma observed in the H&E stain shown in Fig. 2. In the 3-month refed group (r3m), the number of apoptotic nuclei was lower than that observed in the deficient group (d3m) but higher than that in the control group (r3m vs c3m). Moreover, the glandular parenchyma of the r3m group increased considerably compared to the deficient group (d3m) (Fig. 4A).

**Figure 4:**
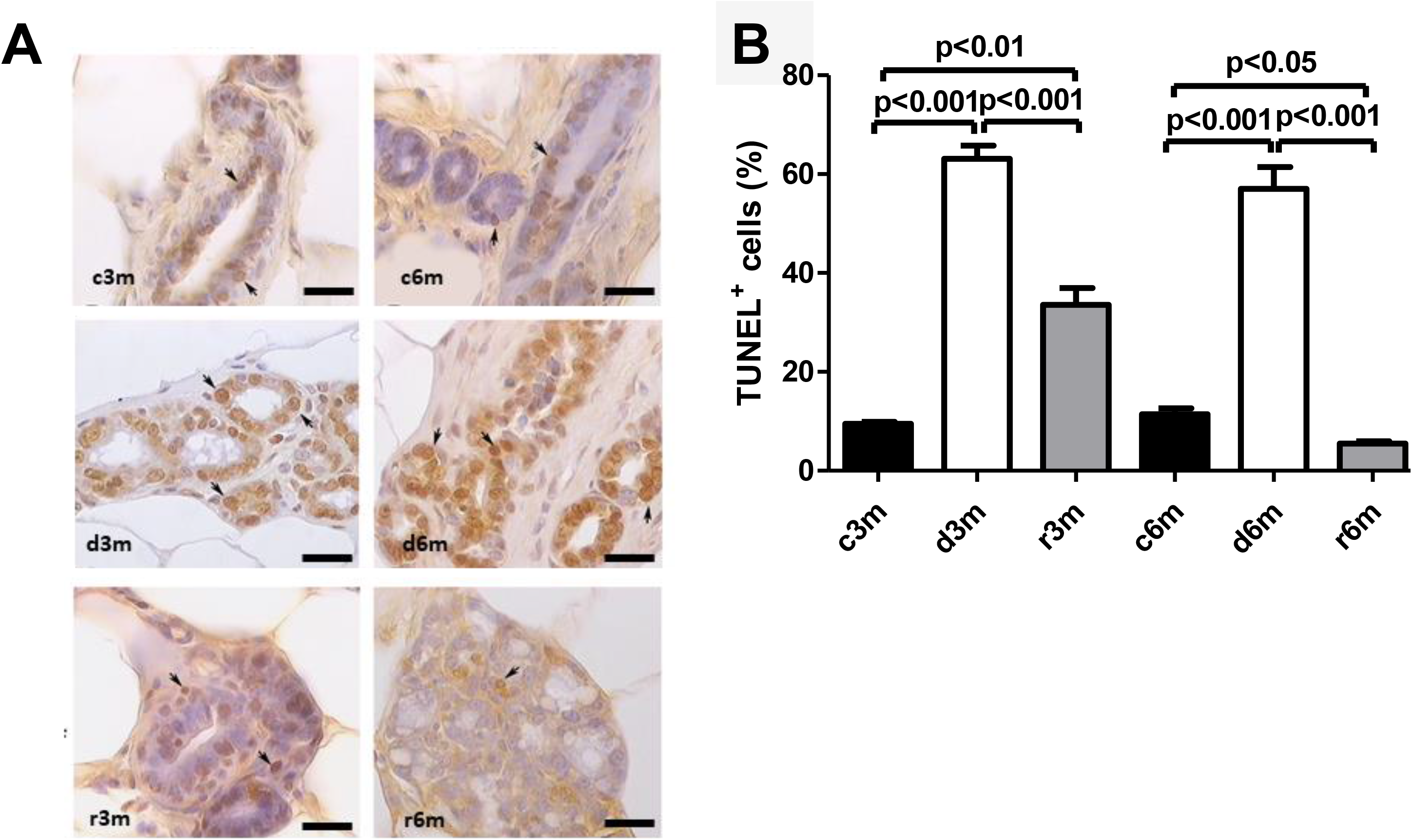
VAD and further refeeding with VAD cause apoptosis changes in the mammary gland. A) TUNEL immunohistochemistry of mammary glands from the different experimental groups shown in Figure 1. Magnification 100×. The scale bar is 25 μm, and a black arrow indicates positive nuclei. B) Percentage of TUNEL-positive cells in the mammary gland. Data depict representative images or mean values ± SDs (n = 4).

The mammary gland of control rats in the 6-month regime has very few cells that present apoptotic nuclei (c6m). At 6 months, a greater number of TUNEL^+^ cells was observed in the deficient group than in the control group (d6m vs c6m). In the c6m group, normal development of the glandular parenchyma, with secretion inside the acini, was observed. The number of TUNEL^+^ cells in the r6m group was lower than that in the mammary gland of the c6m group (Fig. 4A). These data from 3- and 6-month feeding regimes were corroborated by cellular quantification (Fig. 4B).

### 3.6. Subchronic dietary vitamin A deficiency affects cell proliferation in the mammary gland of nulliparous rats

Cell proliferation is an important tissue response to cell death and tissue dysplasia; thus, we measured PCNA to understand how tissue homeostasis is affected by subchronic dietary VAD. Figure 5 shows the results of PCNA immunohistochemistry (Fig. 5A) and the corresponding quantification of PCNA+ cells (Fig. 5B). In the 3-month control group, nuclear immunostaining was emphasized in the alveolus, whereas in the deficient rats at 3 months, a decrease in immunostaining was observed in the control group (d3m vs c3m). In the 3-month refed group, an increase in immunostaining was observed with respect to the deficient group (r3m vs d3m). In the 6-month control group, PCNA immunostaining was emphasized in numerous alveoli and ducts, whereas in the deficient rats of the 6-month group, a decrease in PCNA^+^ cells compared to the control was observed (d6m vs c6m). In the r6m group, there was an increase in PCNA immunostaining compared to that in the d6m group. These observations were corroborated by cellular quantification (Fig. 5B). Dietary vitamin A maintains secretory cell proliferation in the mammary gland of nulliparous rats.

**Figure 5:**
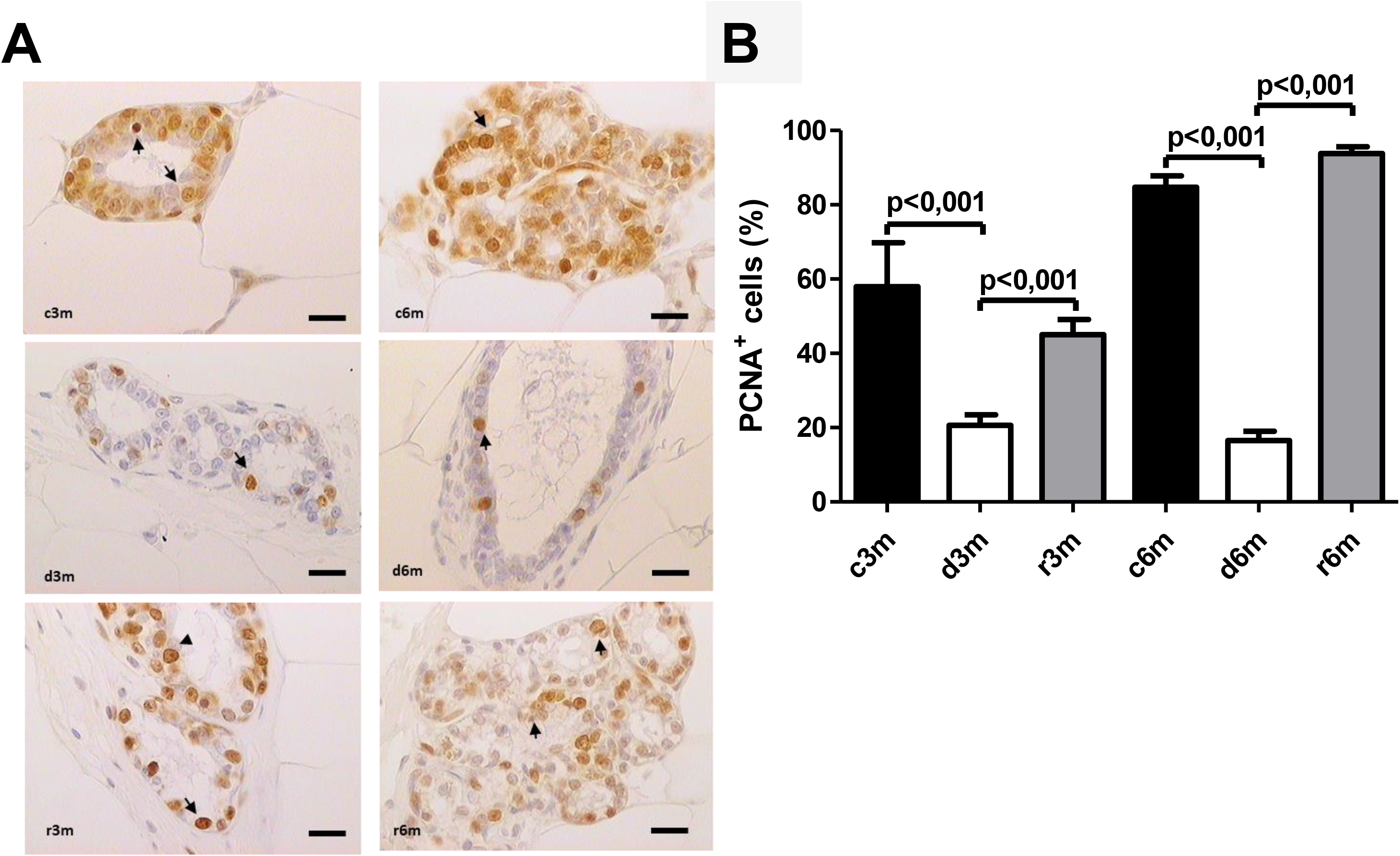
VAD and further refeeding with a VAS diet causes changes in cell proliferation in the mammary gland. A) Immunohistochemistry for PCNA in the mammary gland. Black arrows indicate positive PCNA+ cells. Magnification 100×. The scale bar is 25 μm. B) Quantification of the percentage of PCNA+ cells from the immunohistochemistry images shown in A. Data are shown as representative images or mean values ± SD (n = 10).

### 3.7. Dietary VAD alters the balance between apoptosis and proliferation

The TUNEL/PCNA ratio increased significantly in the mammary glands of the VAD groups (d3m and d6m) with respect to the respective controls (c3m and c6m) and refed groups (r3m and r6m) (Fig. 6). This antiproliferative but proapoptotic pattern was prevented in the mammary glands of the refeeding groups (r3m and r6m). These data are consistent with a strong modulating effect of dietary vitamin A deficiency on apoptosis and proliferation in the mammary gland of nulliparous rats.

**Figure 6:**
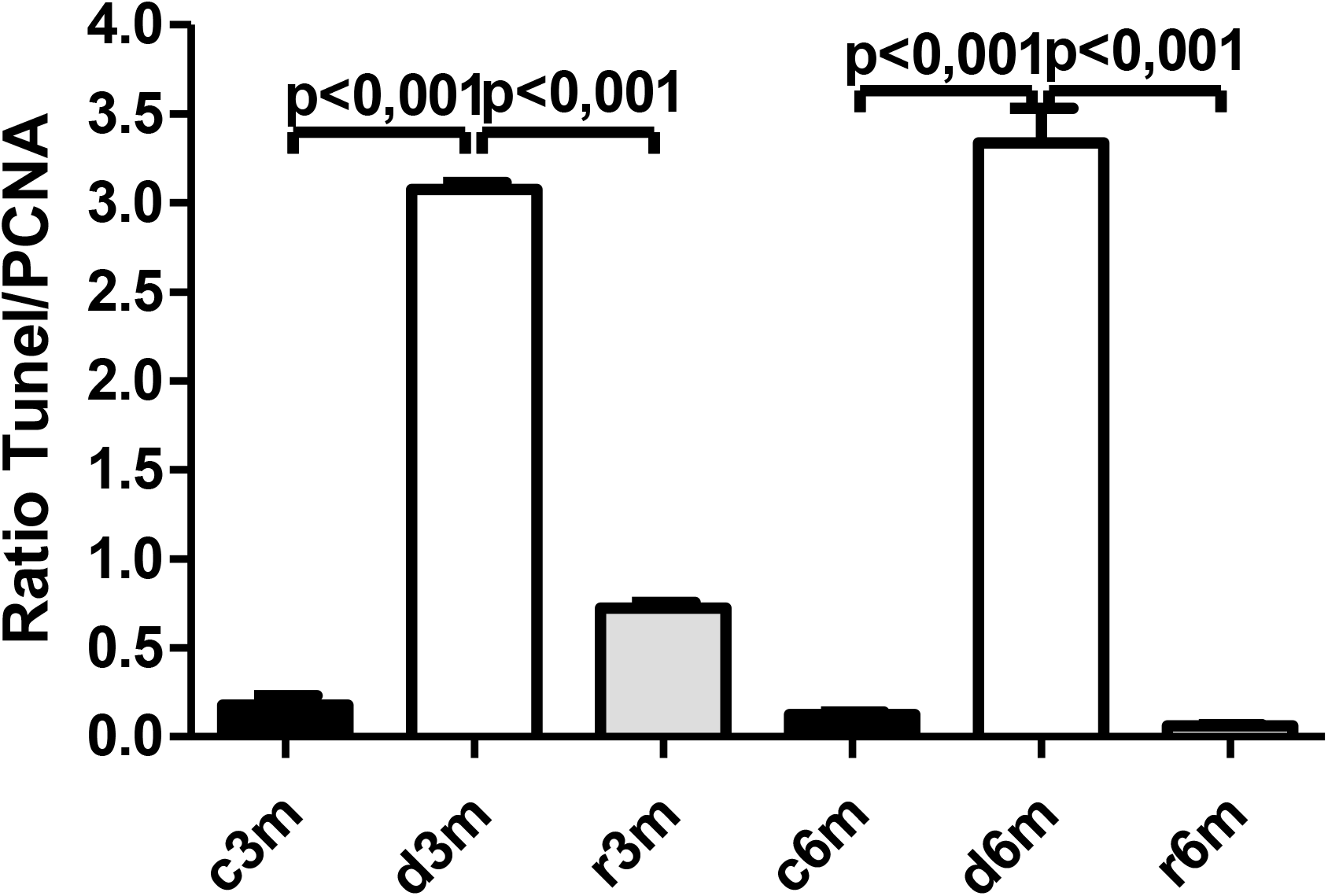
VAS and further refeeding with a VAS diet causes changes in apoptosis and proliferation in the mammary gland. Determination of TUNEL+/PCNA+ cell ratios from the immunohistochemistry images shown in Figs. 5A and 6B. Data depict mean values ± SD (n = 4).

### 3.8. Subchronic dietary deficiency of vitamin A inhibits RAR**α** gene expression in the mammary gland of nulliparous rats

The expression of RARα in the mammary gland of rats from the different experimental groups was determined by RT[qPCR (Fig. 7). Feeding virgin rats a VAD for 3 or 6 months caused reduced RARα gene expression. Refeeding a VAS diet did not reestablish control levels RARα gene expression in the 3-month experimental model (c3m vs r3m). However, one month of refeeding in the 6-month experimental model (r6m) reestablished RARα gene expression to that observed in rats fed a VAS diet for 6 months (c6m).

**FIGURE 7.**
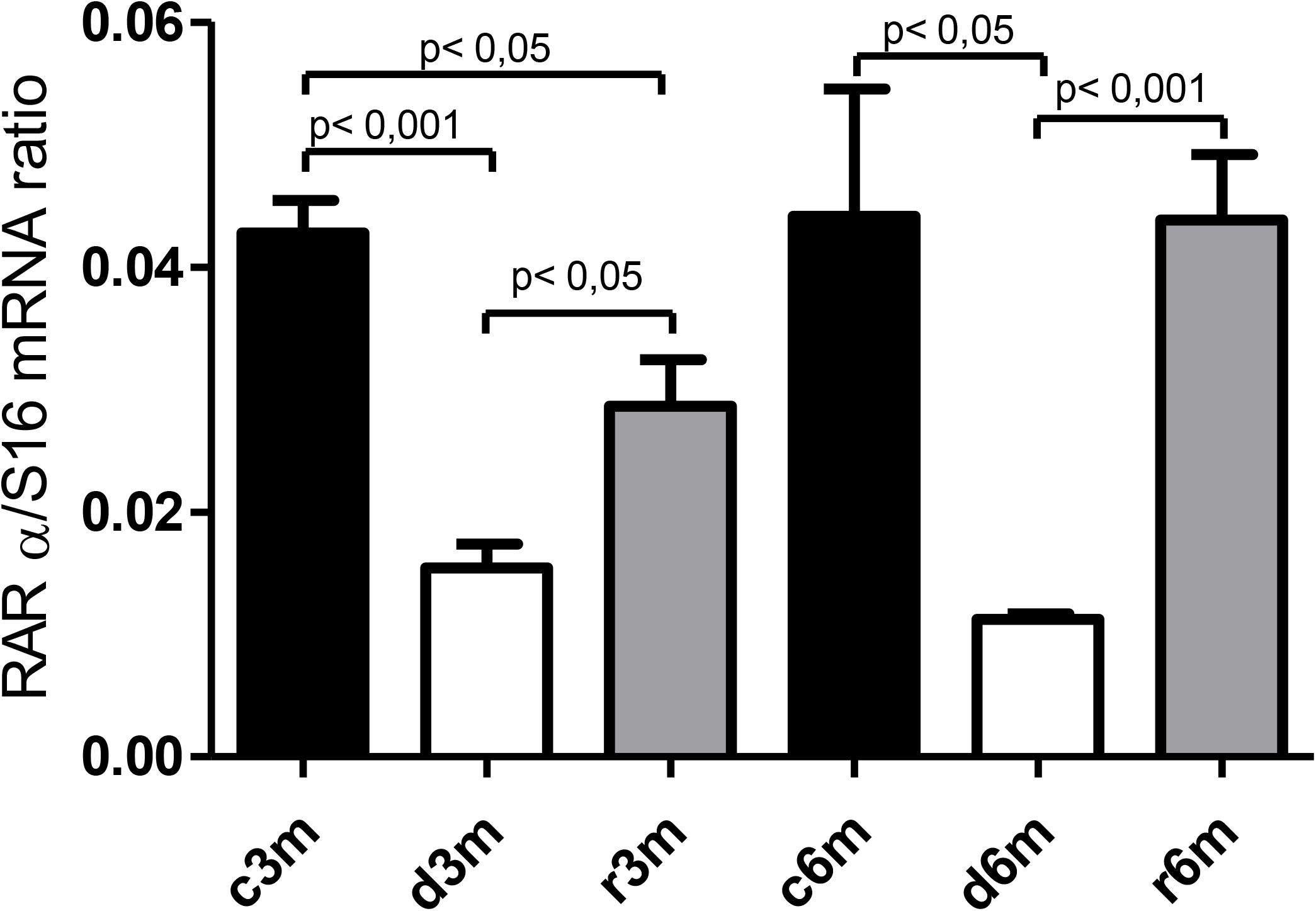
VAD and further refeeding with a VAS diet cause changes in RARα gene expression in the mammary gland. RARα gene expression was assessed by RT[qPCR with respect to S16 gene expression in the different experimental groups. Data are shown as the mean values of the RARα/S16 ratios ± SD (n=4).

### 3.9. Dietary VAD causes inflammation in the mammary gland

Figure 8A shows the TNFα gene expression in the mammary gland in the different experimental groups. At 3 and 6 months of treatment, TNFα expression increased with respect to the control groups. This increase was larger in the d6m group than in the d3m group. This can be explained by the fact that the animals in the 6-month group were older and more prone to inflammatory responses than those in the 3-month group. It may be that younger animals exposed to a VAD diet respond better than older animals (d3m vs r3m and d6m vs r6m). We observed a similar expression pattern to TNFα gene expression when another gene under the transcriptional control of NFκB (e.g., cyclooxygenase2) was analyzed (*data not shown*). Figure 8B shows NFκB gene expression in the mammary gland in the different experimental groups. NFκB gene expression increased in the mammary gland exposed to a VAD diet (c3m vs d3m and c6m vs d6m). In both experimental models, refeeding a VAS diet restored NFκB gene expression to control levels. Dietary vitamin A deficiency causes inflammation in the mammary gland of nulliparous rats.

**Figure 8:**
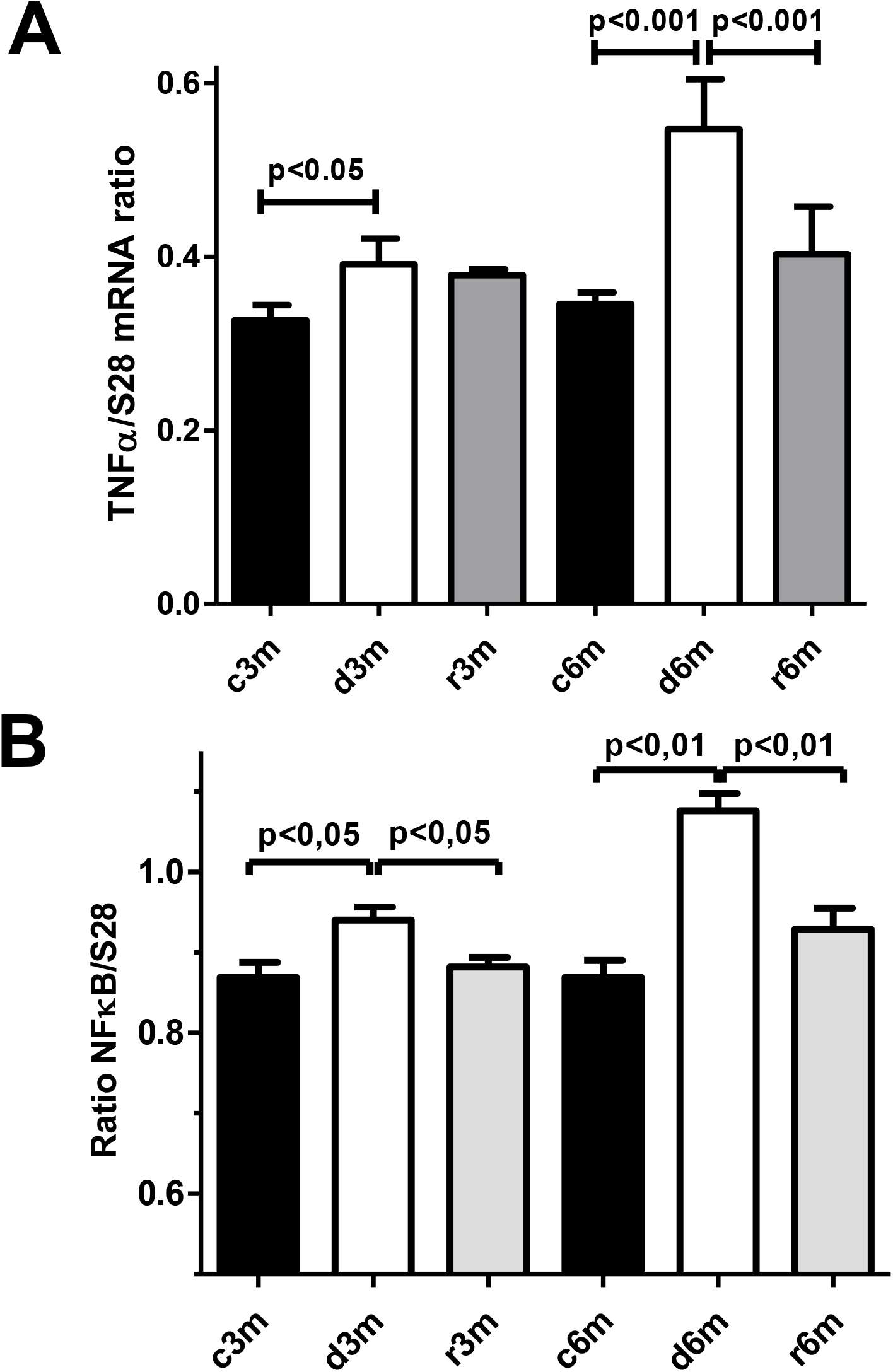
Inflammation response in the mammary gland of rats fed a VAD. A) TNFα gene expression, and B) NFκB gene expression. Gene expression was assessed by RT[qPCR with respect to ribosomal S28 gene expression as a housekeeping gene. The data shown are the mean values of the ratios ± SD (n=8).

## 4. DISCUSSION

Herein, we used two experimental nulliparous-rat models of subchronic dietary vitamin A deficiency (3- and 6-month regimes) and intervention with a VAS diet to understand the mechanism of mammary gland dysfunction as observed in women with dietary deficiency of vitamin A ^1^. Data from our experimental model show that dietary vitamin A is critical for the development of the mammary gland in virgin rats. This development is supported by a tightly regulated balance between the anti-inflammatory, anti-apoptotic, and pro-proliferating effects of dietary vitamin A to support an adequate mammary gland structure and function (Fig. 9). Our exciting data are consistent with an inflammation-induced apoptotic process in the mammary gland of nulliparous rats fed a VAD. Inflammation caused by the absence of RA-RARα mediated inhibition of NFκB signaling may be, among others, a cause for enhanced expression of pro-apoptotic mediators, such as BAX, therefore enhanced apoptosis in the mammary gland of rats fed a VAD.

**Figure 9:**
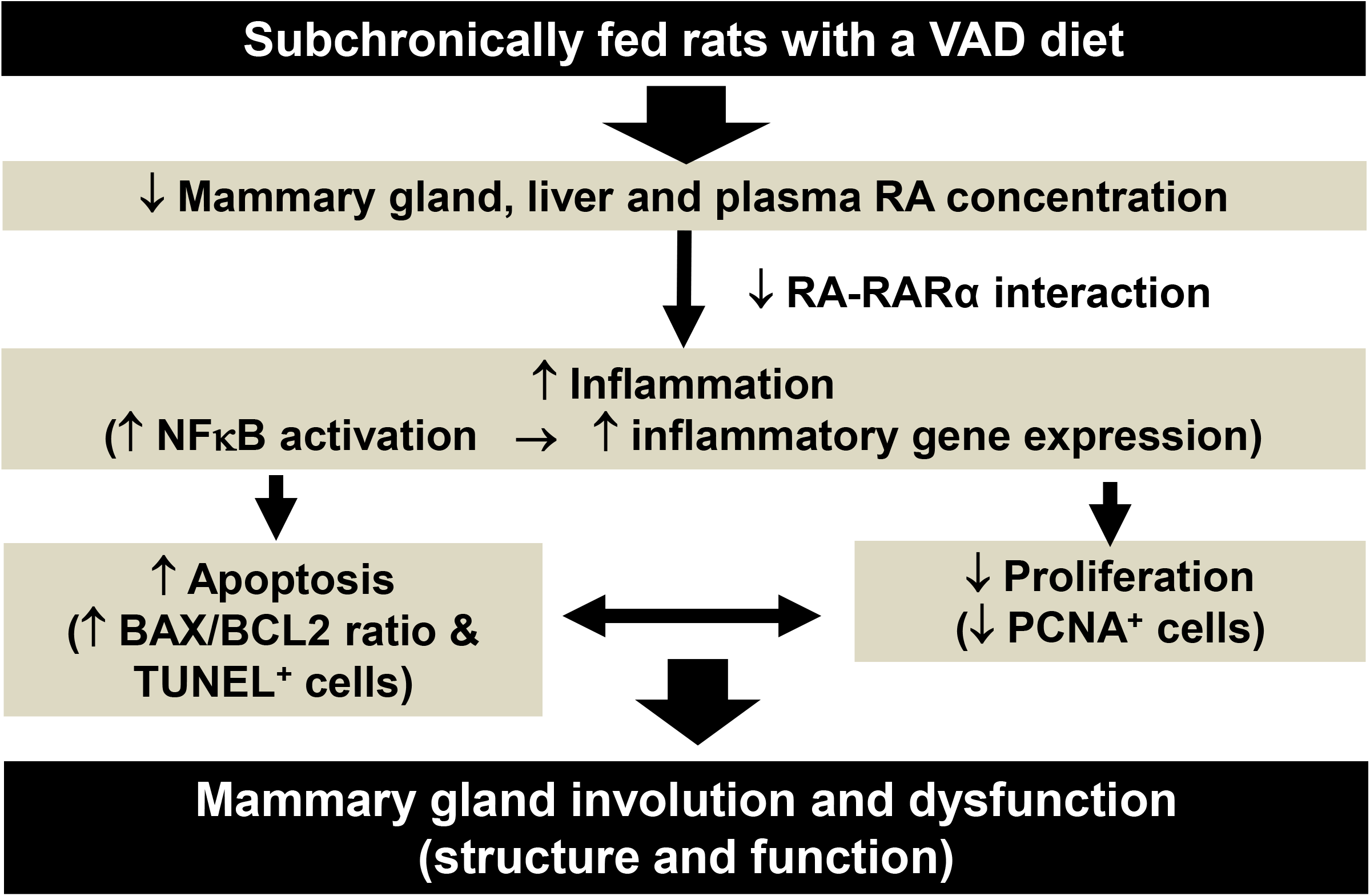
A sub-chronic dietary vitamin A deficiency causes apoptosis, inflammation, and altered cell proliferation in the mammary gland of nulliparous rats. Schematic representation of the main findings of this study. Subchronic dietary VAD reduces tissue and plasma RA concentration. This leads to a reduced activation of the RA-RARα axis causing an enhanced activation of NFκB inflammatory signaling pathway with expression of adhesion molecules for inflammatory cells and pro-inflammatory cytokines. Consequently, mammary gland parenchyma shows inflammatory cell infiltration and apoptosis, but reduced cell proliferation. These changes are consistent with structural and functional alterations causing mammary gland involution and dysfunction. Mammary gland involution and dysfunction are totally or partially reversed by supplementation with a VAS diet.

Vitamin A deficiency is common in the population because it cannot be synthesized by itself in the body and must be actively obtained from food ^1^. Vitamin A is obtained exclusively through dietary sources, primarily in the form of carotenoids and retinyl esters, which can be delivered to peripheral tissues through chylomicron remodeling. Liver and adipose tissue constitute the primary sites of vitamin A storage, with ∼80–85% and ∼15– 20%, respectively, of total retinyl ester and retinol stores in the body ^1^. Bioavailable RA is needed for mediating many physiologically important processes in the body, involving the expression of multiple anti-inflammatory genes and signal transduction pathways^1^.

Bartlett et al. ^30^ reported that circulating insulin-like growth factor-I concentration decreased in vitamin A– deficient rats. It has been reported that VAD leads to reduced body weight in a Japanese quail model ^31^. This may explain the reduced body weight observed in those rats fed a VAD, and serum retinol concentrations are homeostatically controlled and only fall when liver stores of vitamin A are very low ^32^. The RA concentration within tissues is tightly regulated, with liver and adipose tissue being the main stores in the human body ^1^. Vitamin A refeeding leads to further body weight regain in the refeeding groups (r3m and r6m).

Plasma concentrations of RA indicate vitamin A status^1^. Retinol levels are reduced during acute inflammation through increased urinary retinol excretion, decreased gastrointestinal retinol absorption, and lowered synthesis of retinol-binding protein by the liver. Consistent with our data, the serum and tissue concentrations of RA decreased in rats fed a VAD diet for 3 and 6 months. Under our experimental conditions, mammary gland concentrations of RA did not reach control levels when refeeding a VAS diet for 0.5-(r3m) or 1 month (r6m). The largest impact of VAD was found in the older rats exposed to 6 months of VAD diet. This may be explained by the fact that it is more difficult to reach the homeostatic concentration of RA in older rats when refeeding a VAS diet.

Apoptosis is an important process for controlling the growth of normal and neoplastic breast tissues ^33, 34^. BAX protein can suppress the ability of BCL-2 to block apoptosis ^35^. In some tissues, including the breast, stomach, skin, lymph nodes, colon, and small intestine, BAX and BCL2 expression patterns are regulated in parallel, suggesting that there is active antagonism between the two proteins ^36^. Vitamin A deficiency can cause diseases such as decreased immunity, night blindness, dry skin, diarrhea, and certain cancers, including breast cancer^37^. A recent meta-analysis has shown that in North American and Asian women populations, high dietary consumption of vitamin A or supplements decreases the incidence of breast and ovarian cancers ^38^.

Furthermore, RA signaling is needed not only for morphogenesis and development of the gland and adequate milk production but also during the weaning process, when epithelial cell death is coupled with tissue remodeling ^6, 7^. These pieces of evidence are consistent with our findings showing several structural abnormalities found in the mammary tissue from our d3m and d6m experimental groups. These abnormalities are restored by refeeding a VAS diet.

Transcriptional activation of the nuclear receptor RAR/RXR by RA often leads to inhibition of cell growth. However, in some tissues, RA promotes cell survival and hyperplasia, activities that are unlikely to be mediated by RAR. Opposing effects of RA on cell growth emanate from alternate activation of two different nuclear receptors ^39^. Interestingly, our data show that refeeding the rats with a VAS diet restored the anti-apoptotic/pro-apoptotic balance in the mammary gland. Furthermore, BAX gene expression was significantly increased and BCL-2 expression was significantly decreased in PC12 cells transduced with Ad-siRAR-α after oxygen-glucose deprivation-induced injury at the mRNA and protein level ^40^. In agreement with our data, this study suggests that the interaction of RA with its cognate receptor, RARα, inhibits BAX gene expression. Therefore, subchronic deficiency of dietary vitamin A might inhibit this pathway, thus inducing BAX expression and the apoptotic process.

PCNA is a known marker of proliferation that controls mammary gland development. In our 3- and 6-month models of vitamin A deficiency, a significant decrease in PCNA immunostaining was observed with respect to the control and refed groups. RA positively regulates PCNA expression in embryos ^41^. In addition, refeeding with a VAS diet normalizes the cell proliferation marker PCNA and restores mammary gland structure. The *N*-terminal end of PCNA interacts directly with RARE, thus affecting the transcriptional response to RA in a promoter-specific way ^42^. In a model of quail vitamin A deficiency, cell division in the spinal cord, free of RA, decreased in the stages of neural differentiation ^43^. Matthews et al. ^44^ observed a decrease in the cell mass in the pancreas of vitamin A-deficient rats, attributed to a reduction in the rate of cell replication.

Our data show that feeding rats a VAD diet increases markers of inflammation. RA- RAα exerts an anti-inflammatory effect by affecting the NFκB and TNFα signaling pathways ^45, 46^. For instance, a*ll-trans*-retinoic acid inhibits cyclooxygenase-2 and TNFα gene expression induced by bacterial lipopolysaccharide (LPS) in murine peritoneal macrophages ^47^. Moreover, RA treatment inhibits IL-12 production in LPS-activated macrophages. This is caused by the competitive recruitment of transcription integrators between NFκB and RA-RXR for the NF-response elements located in the regulatory region of several pro-inflammatory and pro-apoptotic genes ^45^.

NFκB and TNFα regulate apoptosis and involution of the mammary gland ^1^. It is noteworthy that NFκB is also involved in inflammatory responses, and it is conceivable that these two signaling pathways govern not only the death/survival balance but also the inflammatory response ^33^.

In summary, our data are consistent with vitamin A deficiency leading to a reduced negative modulatory effect of RARα-RA competing for NFκB response elements, which leads to increased inflammation mediators, reduced cell proliferation, and expression of pro-apoptotic markers. RA deficiency leads to abnormalities in mammary gland development in nulliparous rats. These effects of vitamin A deficiency can be reversed by vitamin A dietary supplementation.

## AUTHORS CONTRIBUTIONS

VGM performed the *in vivo* experiments, tissue collection, and all experiments/measurements. FV and MF performed histological analyses, AM performed chemical analyses of food, CVF performed apoptosis measurements, FC performed biochemical analyses, GJ, MSG, DCR, and SEGM coordinated the study design, obtained funding for this study, discussed the data, and wrote the manuscript.

## ACKNOWLEDGMENTS

This research is an IN-LIFE-TRIBUTE to Professor Dr Maria Sofia Gimenez for her highly valuable contribution to the scientific field of nutrition and on the impact of environmental toxicants on metabolism, as well as for her effective career as a mentor in the formation of highly qualified human resources during her academic and scientific career at the National University of San Luis and CONICET.

The authors acknowledge the valuable contribution of Lic. Fabricio Penna (FCH, UNSL) on statistic analyses.

This work has been supported by CONICET (PUE013) and the National University of San Luis (PROICO 8104), Argentina. This research was also supported in part by the PICT- 2018-03435 and PICT-2021-I-A-00147 to DCR and SEGM.

