## Supplementary material for "Vitamin a deficiency causes apoptosis in the mammary gland of rats": SUPPL. TABLES

### SUPPLEMENTARY TABLES

#### TABLE S1: AIN-93G diet composition

| Components | g/kg diet |
|---|---|
| **Casein** | 226.0 |
| **Dextrin** | 132.0 |
| **Sucrose** | 100.0 |
| **Corn oil** | 70.0 |
| **Fiber** | 50.0 |
| **Mineral mixture*** | 35.0 |
| **Vitamin mixture**** | 10.0 |
| **L-cystine** | 3.0 |
| **Choline Bitartrate** | 2.5 |
| **Ascorbic acid** | 0.1 |

(*) The composition of the mineral mixture is shown in Table S2, (**) The composition of the vitamin mixture is shown in Table S3. In the case of the VAD diet the vitamin mixture does not contain *trans*-retinyl palmitate.

#### TABLE S2: Vitamin mixture (AIN-93G)

| Vitamin | g/kg |
|---|---|
| **Nicotinic acid** | 3.000 |
| **Calcium pantothenate** | 1.600 |
| **Pyridoxine-HCl** | 0.700 |
| **Thiamine-HCl** | 0.600 |
| **Riboflavin** | 0.600 |
| **Folic acid** | 0.200 |
| **D-biotin** | 0.020 |
| **Vitamin B-12 (cyanocobalamin)** | 2.500 |
| **Vitamin E (500 IU/g)** | 15.000 |
| **Vitamin A (*trans*-retinylpalmitate)[&]** | 0.800 |
| **Vitamin D3 (400,000 IU/g)** | 0.250 |
| **Vitamin k** | 0.075 |
| **Sucrose** | 974.655 |

(**&**) The vitamin mixture added to the VAS and VAD have the same composition except that in the VAD diet the vitamin mixture does not contain *trans*-retinyl palmitate.

#### TABLE S3. Mineral mixture (AIN-93G)

| Minerals | mg/kg mixture |
|---|---|
| ***(a)* Essential mineral elements:** | |
| Calcium carbonate, anhydrous | 357.00 |

| | |
|---|---|
| Potassium phosphate, monobasic | 196.00 |
| Potassium Citrate, Monohydrate | 70.78 |
| Sodium chloride | 74.00 |
| Potassium sulfate | 46.60 |
| Magnesium oxide | 24.00 |
| Ferric citrate | 6.06 |
| Zinc carbonate | 1.65 |
| Manganese carbonate | 0.63 |
| Cupric carbonate | 0.30 |
| Potassium iodide | 0.01 |
| Sodium selenate, anhydrous | 0.01025 |
| Ammonium paramolybdate.4$H_2O$ | 0.00795 |
| ***(b)* Potentially beneficial elements:** | |
| Sodium meta-silicate.9$H_2O$ | 1.4500 |
| Chromium potassium sulfate.12$H_2O$ | 0.2750 |
| Lithium chloride | 0.0174 |
| Boric acid | 0.0815 |
| Sodium fluoride | 0.0635 |
| Nickel carbonate | 0.0318 |
| Ammonium vanadate | 0.0066 |
| Sucrose | 221.0260 |

### Table S4: Sequences of primers used in RT-qPCR measurements

| Gen | Forward (5'- 3') | Reverse (5'- 3') | Gen Bank Access |
|---|---|---|---|
| ***rarα*** | CGCCTGTGAGGGCTGTAAG | ATGCCCACTTCGAAGCATTT | NM_031528 |
| ***bcl2*** | TGGATGACTGAGTACCTGAAC | AGAGACAGCCAGGAGAAATCAAAC | NM_016993.1 |
| ***bax*** | TGGTTGCCCTTTTCTACTTTGC | TGATCAGCTCGGGCACTTTA | NM_017059.2 |
| ***nfκb*** | AGCAACCGAAACAGAGAGG | TTTGCAAAGCCAACCACCAT | NM001276711.1 |
| ***tnfα*** | GGTGATCGGTCCCAACAAGGA | CACGCTGGCTCAGCCACT | NM012675.3 |
| ***S28*** | GTGAAAGCGGGGCCTCACGATCC | GTACTGAGCAGGATTACCATGGC | NR046239.1 |
| ***S16*** | TCCAAGGGTCCGCTGCAGTC | CGTTCACCTTGATGAGCCCATT | NM_001169146.1 |
